# SapTrap assembly of *C. elegans* MosSCI transgene vectors

**DOI:** 10.1101/805507

**Authors:** Xintao Fan, Sasha De Henau, Julia Feinstein, Stephanie I. Miller, Bingjie Han, Christian Frøkjær-Jensen, Erik E. Griffin

**Affiliations:** Department of Biological Sciences, Dartmouth College, Hanover NH 03755; Center for Molecular Medicine, Molecular Cancer Research, University Medical Center Utrecht, 3584 CG Utrecht, The Netherlands; King Abdullah University of Science and Technology (KAUST), Biological and Environmental Science and Engineering Division (BESE), KAUST Environmental Epigenetics Program (KEEP), Thuwal 23955-6900, Saudi Arabia

**Keywords:** MosSCI, *C. elegans*, mitochondria, endoplasmic reticulum, SapTrap

## Abstract

The Mos1-mediated Single-Copy Insertion (MosSCI) method is widely used to establish stable *Caenorhabditis elegans* transgenic strains. Cloning MosSCI targeting plasmids can be cumbersome because it requires assembling multiple genetic elements including a promoter, a 3’UTR and gene fragments. Recently, Schwartz and Jorgensen developed the SapTrap method for the one-step assembly of plasmids containing components of the CRISPR/Cas9 system for *C. elegans* (Schwartz and Jorgensen 2016 Genetics, 202:1277-1288). Here, we report on the adaptation of the SapTrap method for the efficient and modular assembly of a promoter, 3’UTR and either 2 or 3 gene fragments in a MosSCI targeting vector in a single reaction. We generated a toolkit that includes several fluorescent tags, components of the ePDZ/LOV optogenetic system and regulatory elements that control gene expression in the *C. elegans* germline. As a proof of principle, we generated a collection of strains that fluorescently label the endoplasmic reticulum and mitochondria in the hermaphrodite germline and that enable the light-stimulated recruitment of mitochondria to centrosomes in the one-cell worm embryo. The method described here offers a flexible and efficient method for assembly of custom MosSCI targeting vectors.

## Introduction

The rich toolbox of techniques available to manipulate gene expression in *C. elegans* is a major attraction of this model organism. Several approaches have been developed to introduce transgenes and to induce efficient CRISPR/Cas9 mediated gene editing (Nance and Frøkjær-Jensen, 2019). The Mos1-mediated Single-Copy Insertion (MosSCI) method has been widely adopted to introduce transgenes in *C. elegans* because single-copy transgenes are integrated at defined chromosomal positions, thereby mitigating potential concerns of transgene integration at random positions (Frøkjær-Jensen et al., 2012; Frøkjær-Jensen et al., 2008; Frøkjær-Jensen et al., 2014). MosSCI transgene integration results from homologous recombination between a MosSCI targeting vector containing the transgene construct and one of the safe-harbor integration sites that have been engineered at defined positions in the genome.

Transgenes typically include multiple genetic elements including a promoter, one or more gene fragments and a 3’UTR. A number of strategies can be used to assemble these elements together including traditional restriction enzyme cloning, Gateway cloning (Hartley et al., 2000), *in vivo* recombineering (Philip et al., 2019) or Gibson cloning (Gibson et al., 2009). Each of these strategies has both advantages and disadvantages. For example, Gateway cloning allows the efficient modular “mix and match” cloning of large collections of promoter, ORF and 3’UTR cassettes (Brasch et al., 2004; Dupuy et al., 2004; Mangone et al., 2010; Zeiser et al., 2011). However, Gateway cloning can be expensive due to the required use of proprietary enzyme mixes and leaves ~25 base pair *att* recombination site “scars” at each cassette junction. In contrast, Gibson cloning allows the efficient, “scar-free” assembly of multiple gene fragments but does not allow the “mix and match” cloning of existing cassettes, making this approach laborious if many constructs are needed.

Schwartz and Jorgensen recently developed the SapTrap method for efficient, modular and single step assembly of CRISPR/Cas9 vectors for *C. elegans* (Schwartz and Jorgensen, 2016). The SapTrap method is based on the Golden Gate cloning technique (Engler et al., 2008) and takes advantage of the SapI type II restriction enzyme, which cuts DNA at defined positions adjacent to its recognition sequence to generate three-base 5’ overhangs. By designing SapI restriction fragments with complementary overhangs, multiple fragments can be assembled together in a defined order in a single digestion and ligation reaction. In this study, we report on the adaptation of the SapTrap system for the efficient, inexpensive, modular, and “scar-free” assembly of transgenes in a MosSCI targeting vector. We have developed a toolkit for expression of transgenes in the *C. elegans* germline, including a collection of cassettes containing tags for fluorescence imaging and for the ePDZ/LOV optogenetic system (Fielmich et al., 2018; Strickland et al., 2012). As a proof of principle, we have used this system to generate a collection of mitochondrial and endoplasmic reticulum reporter strains and a strain in which light induces the transport of mitochondria to centrosomes in the one-cell worm embryo.

## Results and Discussion

### Adaptation of the SapTrap system for cloning MosSCI targeting vectors

To adapt the SapTrap approach (Schwartz and Jorgensen, 2016) for the assembly of MosSCI targeting vectors, we started by making two changes to the universal MosSCI targeting vector pCFJ350 (Frøkjær-Jensen et al., 2012), which targets transgenes for insertion at the commonly used *ttTi5605* site (Frøkjær-Jensen et al., 2008). First, we introduced single base pair changes to disrupt the two SapI restriction sites located in the “Left” and “Right” homology arms of pCFJ350. Second, we inserted two SapI sites into the multiple cloning site that were oriented such that they are removed from the vector backbone by digestion with SapI. The resulting MosSCI targeting vector was named pXF87 (Figure 1A).

**Figure 1.**
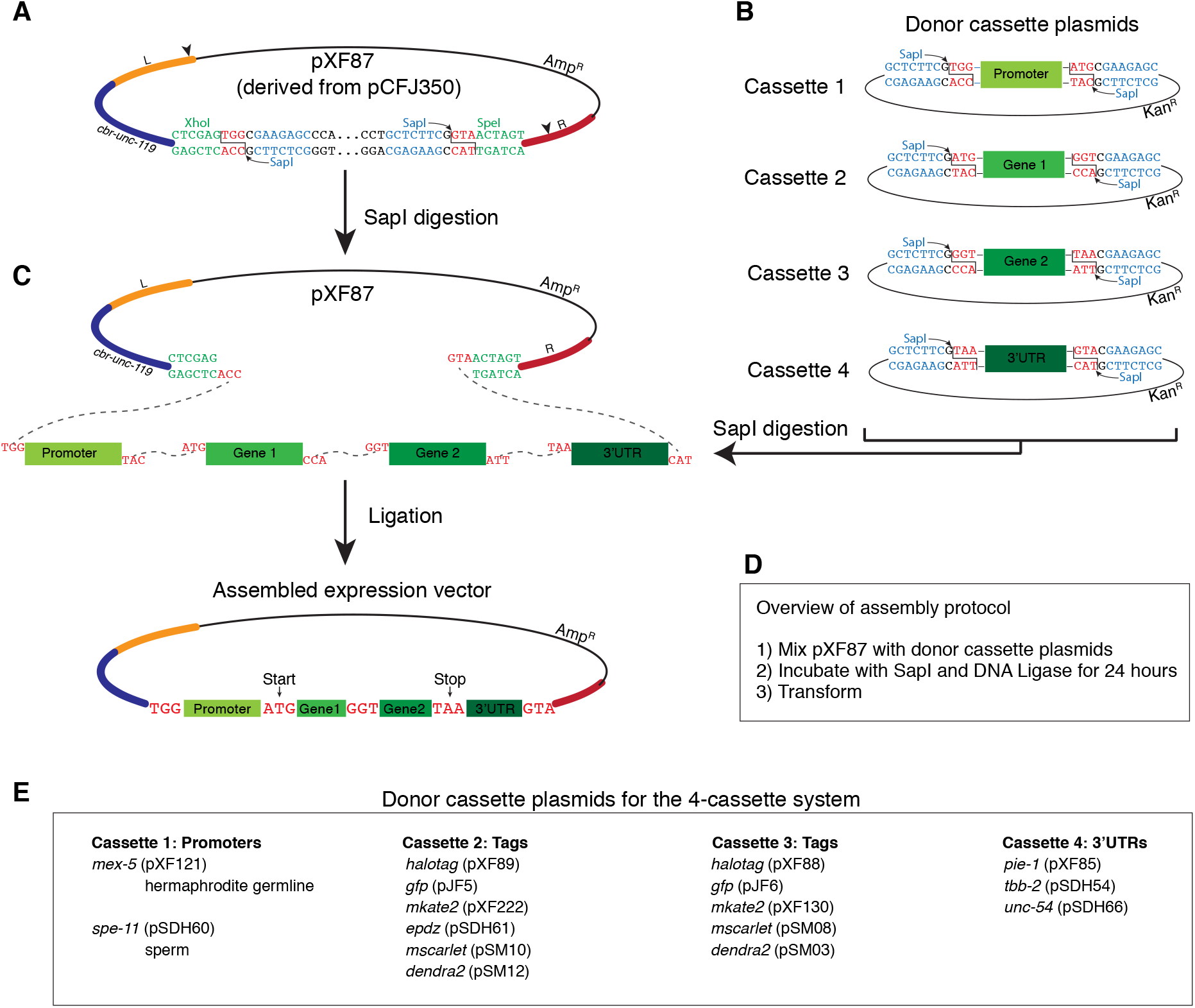
SapTrap assembly of MosSCI targeting vectors using the four-cassette system. **A.** The MosSCI targeting vector pXF87 was derived from pCFJ350 by mutating two SapI restriction sites (indicated by arrowheads in the “Left” (L) and “Right” (R) homology arms) and introducing two SapI sites (blue text) between the XhoI and SpeI sites (green text). SapI cleavage sites are in red text. The SapI recognition sites are oriented such that upon digestion they are removed from the vector backbone. The cbr-*unc-119* gene is used as a positive selection marker to facilitate the identification of transgenic animals. **B.** Design of the donor cassette vectors used for the 4- cassette cloning strategy. **C.** The curved dotted lines indicate the overhangs that anneal during the ligation reaction. **D.** Overview of the assembly protocol. For a detailed protocol, see the Materials and Methods section. **E.** Summary of available promoter, gene tag and 3’UTR donor cassette plasmids.

We next cloned a series of plasmids that contain donor cassettes flanked by SapI restrictions sites (Figure 1B). Following digestion with SapI, the cassettes are liberated from the vector backbone and are flanked by 5’ overhangs that direct their order of assembly in pXF87 (Figure 1C). A four-insert cassette system was designed with a promoter in cassette 1, gene fragments in cassettes 2 and 3 (typically a gene and a tag) and a 3’UTR in cassette 4. To minimize the inclusion of extraneous sequences, the junctions between the first and second cassette is the translation start (ATG), between second and third cassettes is glycine (GGT) and between the third and fourth cassettes is the ochre translation stop codon (TAA) (Figure 1C). Donor cassettes encoding tags (such as fluorescent proteins) include short flexible linkers at the protein fusion site (the carboxy terminus of cassette 2 and the amino terminus of cassette 3) (Supplemental Figure S1- S7). The currently available promoter, tag and 3’UTR donor cassette plasmids are listed in Figure 1E and Table 1.

**Table 1.** Donor cassette plasmids used in this study.

| Name | Description |
| --- | --- |
| **Cassette 1 for 4-cassette or 5 cassette system (5'-TGG ..... 3'-TAC)** |  |
| pXF121 | *mex-5* promoter |
| pSDH60 | *spe-11* promoter |
| **Cassette 2 for 4-cassette or 5-cassette system (5'-ATG-3' ..... 3'-CCA-5')** |  |
| **Tags** |  |
| pXF89 | *halotag* (no STOP codon, PATC introns) |
| pJF5 | *gfp* (no STOP codon, PATC introns) |
| pXF222 | *mkate2* (no STOP codon) |
| pSDH61 | *epdz* (no STOP codon) |
| pSM10 | *mscarlet* (no STOP codon) |
| pSM12 | *dendra2* (no STOP codon) |
| **Genes** |  |
| pJF7 | *tomm-20* (no STOP codon) |
| pSDH50 | *tomm-20* (aa1-55) (no STOP codon) |
| pXF262 | *cox-4* (no STOP codon) |
| pXF250 | *npp-20* (no STOP codon) |
| **Cassette 3 for 4-cassette system (5'-GGT-3' ..... 3'-ATT-5')** |  |
| **Tags** |  |
| pXF88 | *halotag* (includes STOP codon, PATC introns) |
| pJF6 | *gfp* (includes STOP codon, PATC introns) |
| pXF130 | *mkate2* (includes STOP codon) |
| pSM08 | *mscarlet* (includes STOP codon) |
| pSM03 | *dendra2* (includes STOP codon) |
| **ORFs** |  |
| pXF90 | *halotag::HDEL* (includes STOP codon, PATC introns) |
| **Cassette 3A for 5-cassette system (5'-GGT-3' ..... 3'-TGC-5')** |  |
| **Tags** |  |
| pSDH51 | *halotag* (no STOP codon, PATC introns) |
| pSM04 | *mkate2* (no STOP codon) |
| pSDH57 | *mscarlet* (no STOP codon) |
| **Cassette 3B for 5-cassette system (5'-ACG-3' ..... 3'-ATT-5')** |  |
| **Tags** |  |
| pXF276 | *lov* domain (includes STOP codon) |
| pSDH52 | *epdz* (includes STOP codon) |
| pSM05 | *mkate2* (includes STOP codon) |
| **Cassette 4 for 4-cassette or 5-cassette system (5'-TAA-3' ..... 3'-CAT-5')** |  |
| pXF85 | *pie-1* 3'UTR |
| pSDH54 | *tbb-2* 3'UTR |
| pSDH66 | *unc-54* 3'UTR |

Note: For the expression plasmid pXF108, the annealed oligos were used to generate HSP-70 in cassette 2.

The *C. elegans* germline is a notoriously difficult tissue in which to achieve stable transgene expression due to silencing of multi-copy extra-chromosomal arrays (Kelly et al., 1997), single-copy insertions generated by MosSCI (*e.g.*, (Frøkjær-Jensen et al., 2016; Shirayama et al., 2012)) or endogenous genes tagged using CRISPR/Cas9 gene editing (*e.g.*, (Fielmich et al., 2018)). Each of our tag donor cassettes encoding gene tags incorporates at least one modification that buffers against silencing, including the inclusion of PATC introns in HaloTag and ceGFP (Frøkjær-Jensen et al., 2016), the elimination of piRNA binding sites in mScarlet, mKate2 and Dendra2 (Seth et al., 2018; Zhang et al., 2018) and the use of sequence motifs found in native germline genes in ePDZ and the LOV domain (Fielmich et al., 2018).

Similar to the SapTrap method developed by Schwartz and Jorgensen (Schwartz and Jorgensen, 2016), MosSCI targeting vectors were assembled in a single tube by incubating pXF87, four donor cassette plasmids, SapI enzyme, ATP and T4 DNA ligase at 25°C for 22 - 24 hours (Figure 1D and Materials and Methods). This reaction was then transformed into *E. coli* and plasmid clones were screened by restriction enzyme digestion followed by sequencing. We assembled nine vectors using the 4-cassette system and 32 of 46 (69.6%) of the plasmids screened had the correct restriction digest pattern (Table 2). Of the vectors with the correct restriction digest pattern, 22 of 23 were correct based on Sanger sequencing analysis. Therefore, the SapTrap method provides an efficient method for the assembly of MosSCI targeting vectors.

**Table 2.**
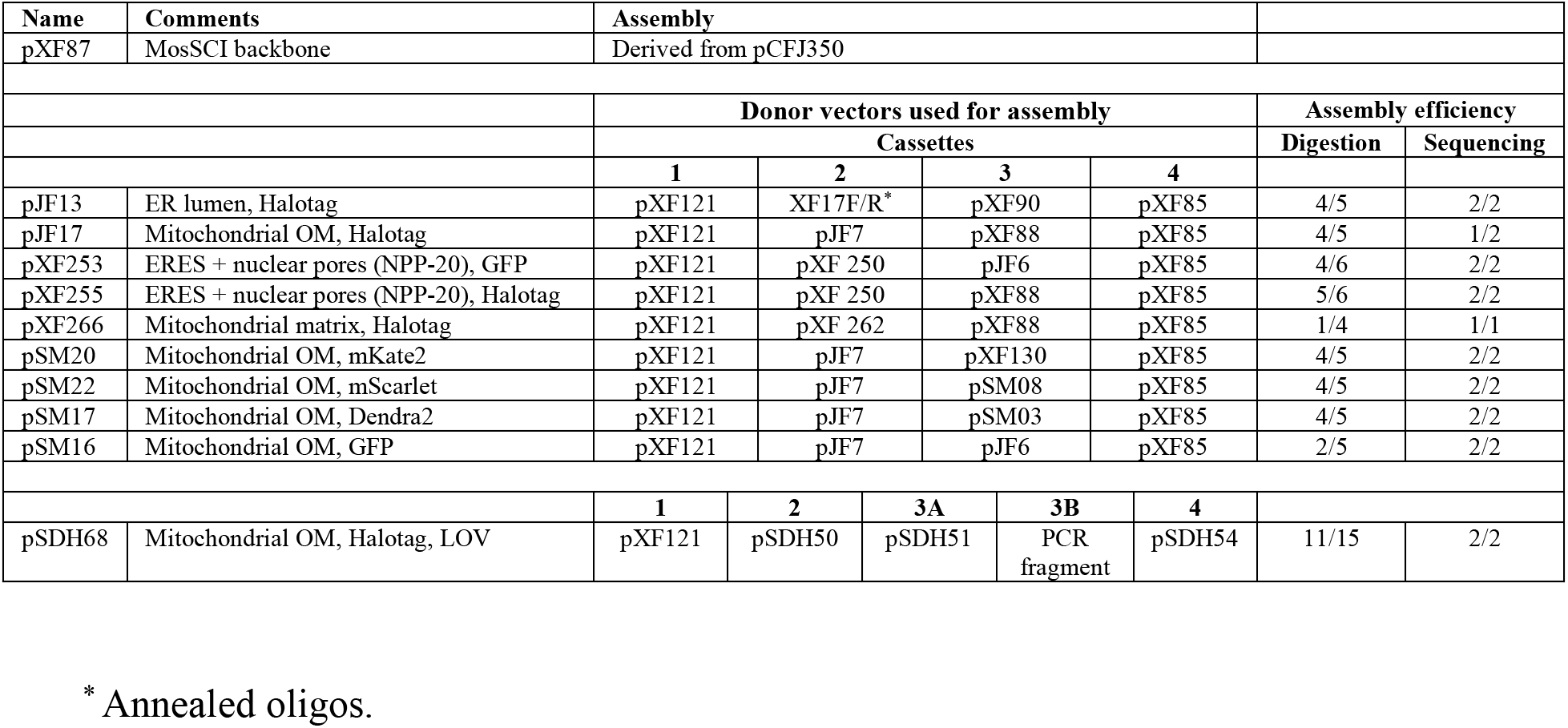
MosSCI targeting vectors used in this study.

### A collection of fluorescent ER and mitochondria strains

We used SapTrap-assembled MosSCI targeting vectors to generate a collection of transgenic strains for analysis of endoplasmic reticulum and mitochondrial dynamics. We first targeted GFP, mKate2, mScarlet, Dendra2 and HaloTag to the cytoplasmic face of the mitochondrial outer membrane by fusing them to the carboxy terminus of TOMM-20. The expression of these transgenes was controlled by the *mex-5* promoter and by the *pie-1* 3’UTR, which results in germline expression that increases around the bend of the adult hermaphrodite gonad (Merritt et al., 2008) (Figure 2A). Strains expressing TOMM-20 fused to HaloTag were labeled with the fluorescent JF_646_ HaloTag ligand (Grimm et al., 2015) by feeding hermaphrodites bacteria mixed with the ligand. Each TOMM-20 fusion protein exhibited the expected tubular localization pattern in the early embryo (Figure 2B-I). We confirmed that TOMM-20::HaloTag colocalized to the same organelle as the mitochondrial matrix protein COX-4::GFP (Raiders et al., 2018) (Figure 2C). We additionally generated strains in which the HaloTag was targeted to the mitochondrial matrix (COX-4::HaloTag) (Figure 2J) and the lumen of the endoplasmic reticulum (HSP-70(aa1-19)::HaloTag::HDEL) (Figure 2K). We fused both GFP and HaloTag to NPP-20, the worm homologue of SEC13, which is both a component of the COPII coat that concentrates to ER exit sites (ERES) (D’Arcangelo et al., 2013) and a component of nuclear pore complexes (Siniossoglou et al., 1996) (Figure 2L, M).

**Figure 2.**
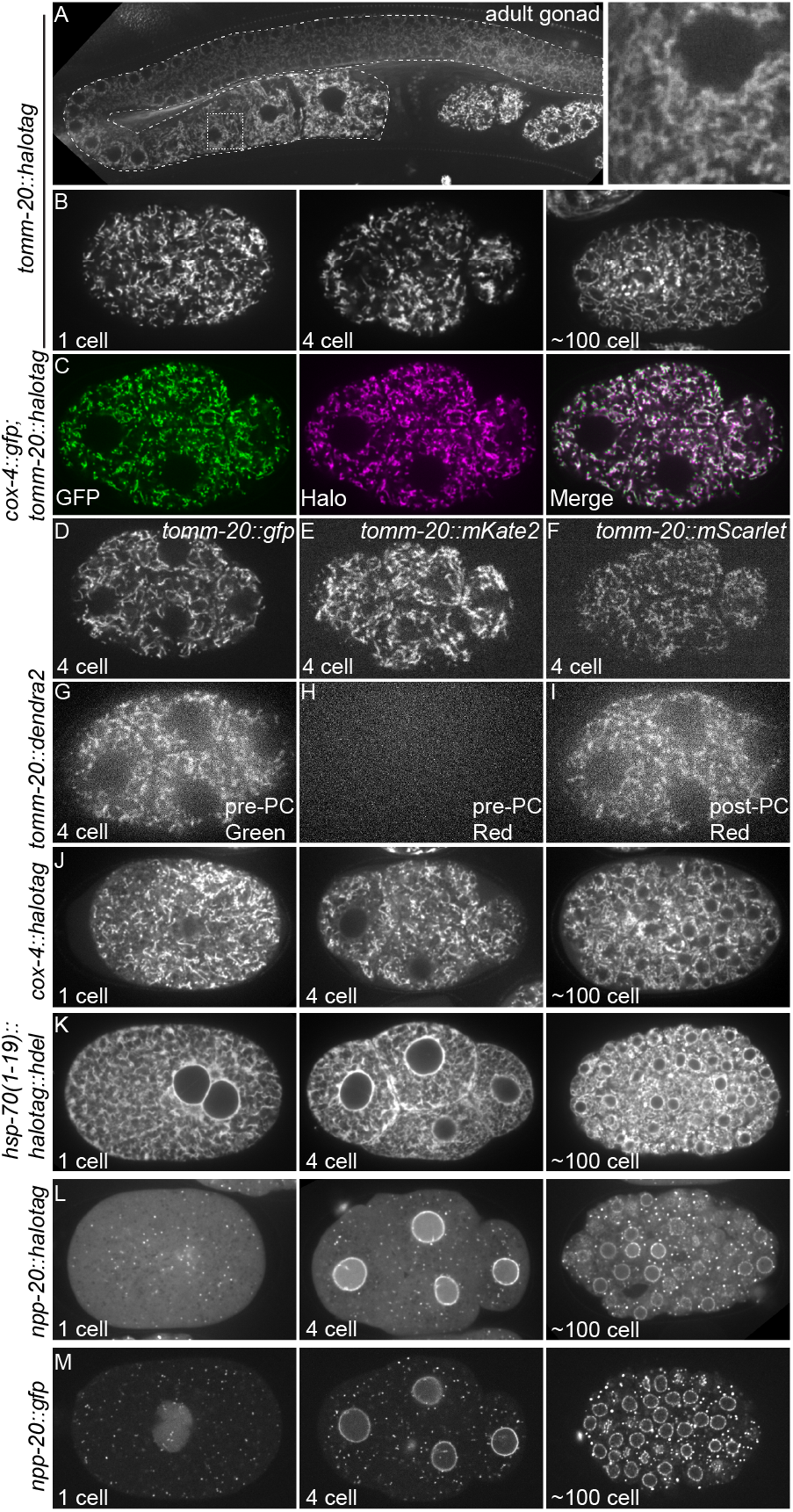
Images of transgenic strains. **A.** Images of TOMM-20::HaloTag labeled with JF_646_ HaloTag ligand in the adult gonad (outlined with curved dotted line), including an inset of the region in the stippled box. **B.** Images of embryos expressing TOMM-20::HaloTag labeled with JF_646_ HaloTag ligand at the 1-cell, 4 cell and ~100 cell stages. **C.** Images of a 4 cell embryo expressing TOMM-20::HaloTag labeled with JF_646_ HaloTag ligand (magenta) and COX-4::GFP (green) (Raiders et al., 2018). **D – F.** Images of embryos expressing the indicated transgenes at the 4-cell stage. **G – I.** Images of a 4 cell embryo expressing TOMM-20::Dendra2 before and after photoconversion (PC). Dendra2 switches from green to red fluorescence upon photoconversion. **J – M.** Images of embryos expressing the indicated transgenes at the 1-cell, 4 cell and ~100 cell stages.

### Five-cassette system

One of the advantages of the SapTrap approach is that it can be easily expanded to include additional insert fragments to create more complex transgenes. To establish a five-cassette system, we used the cassettes 1, 2 and 4 from the four-cassette system and replaced cassette 3 with cassettes 3A and 3B (Figure 3A). We used this approach to generate an optogenetic system to control the localization of mitochondria in the early embryo based on the light induced interaction between the ePDZ and LOV domains (Fielmich et al., 2018; Strickland et al., 2012). We assembled a MosSCI targeting vector that directed expression of TOMM- 20::HaloTag::LOV, which targets the LOV domain to the mitochondrial outer membrane. 11 of 15 assembled plasmids had the corrected restriction digest pattern and 2 of 2 of these plasmids were correct by sequence analysis. A TOMM-20::HaloTag::LOV strain was crossed with a strain in which the dynein heavy chain DHC-1 was fused to ePDZ (Fielmich et al., 2018). Whereas mitochondria in wild-type embryos are dispersed through the cytoplasm (Figure 4A), upon the recruitment of ePDZ::mCherry::DHC-1 to mitochondria by stimulation with 488 nm light, mitochondria were transported on to centrosomes, leaving the peripheral cytoplasm largely devoid of mitochondria (Figure 4B).

**Figure 3.**
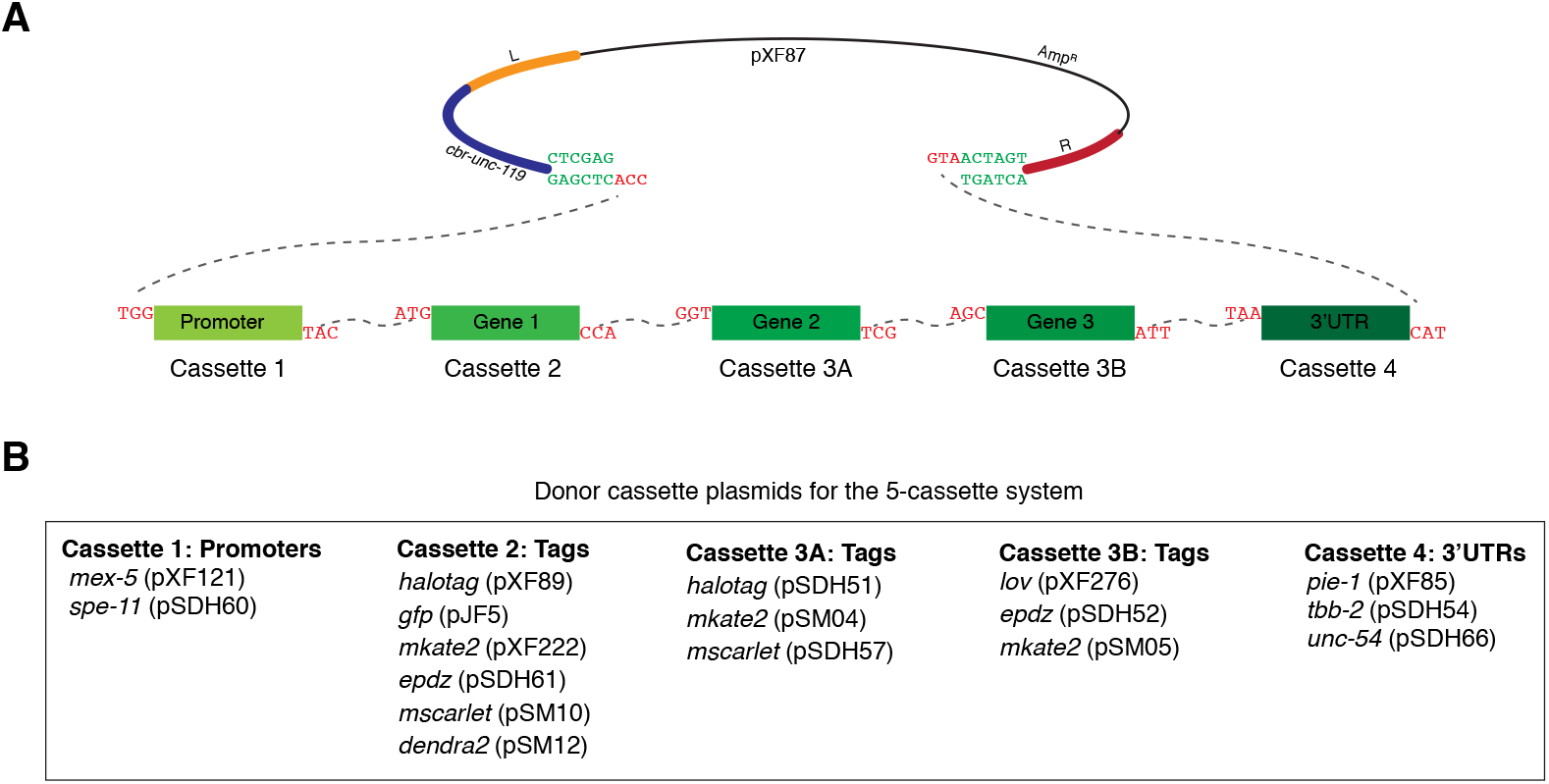
SapTrap assembly of MosSCI targeting vectors using the five-cassette system. **A.** Schematic of pXF87 and the donor cassettes following SapI digestion. The dotted lines indicate the overhangs that anneal during ligation. **B.** Summary of available promoter, gene tag and 3’UTR donor cassette plasmids for the five-cassette system.

**Figure 4.**
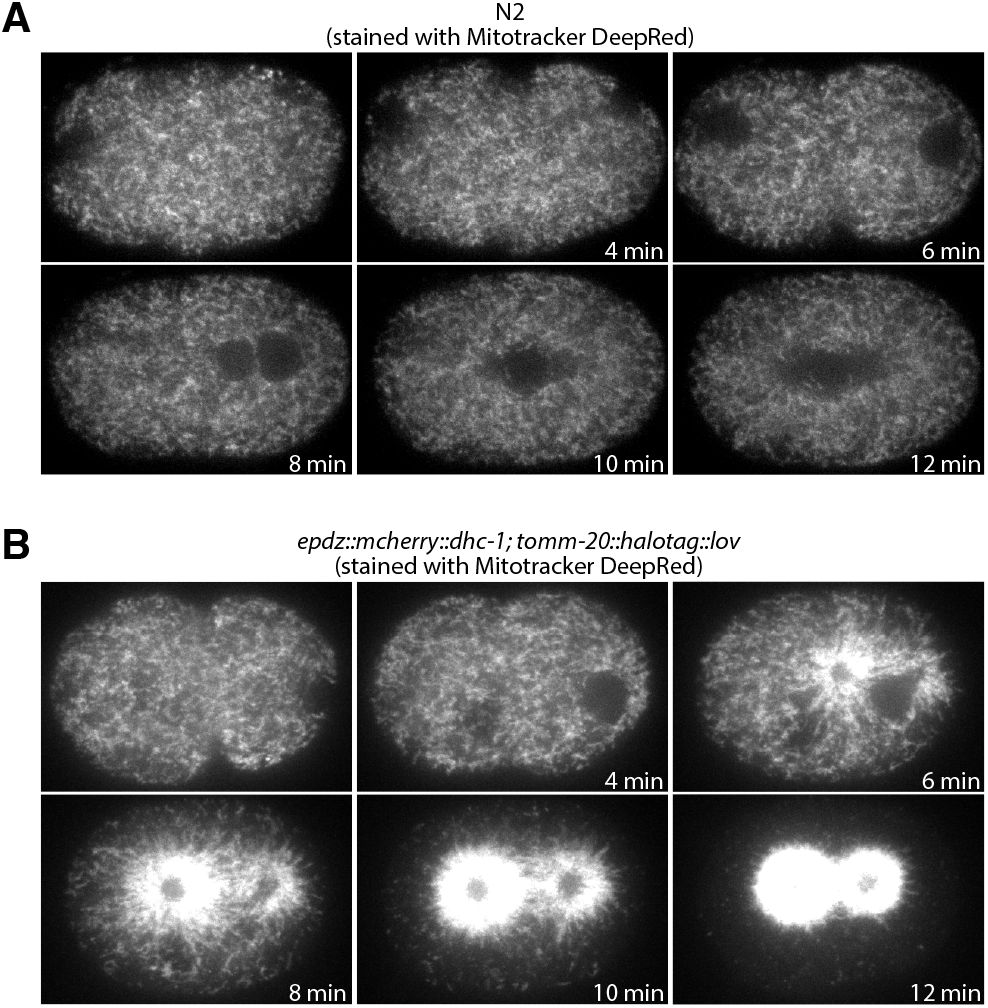
Optogenetic control of mitochondrial distribution in the 1-cell embryo. **A.** Control embryo stained with Mitotracker DeepRed and imaged with 488 nm and 640 nm illumination (640 nm channel shown). **B.** 1-cell *epdz::mcherry::dhc-1*; *tomm-20::halotag::lov* embryo stained with Mitotracker DeepRed and imaged with 488 nm and 640 nm illumination (640 nm channel shown). The 488 nm illumination was used to stimulate the interaction between the ePDZ and LOV domains.

The SapTrap system described here provides an efficient and simple method for the assembly of MosSCI targeting vectors. This approach is similar to the Gateway assembly system (ThermoFisher Scientific) in that once donor cassette plasmids are cloned, they can be assembled in any modular combination. The Gateway system has been widely used to generate MosSCI transgenes and is attractive because there are large collections of promoter, ORF, and 3’UTR donor plasmids available (Brasch et al., 2004; Dupuy et al., 2004; Mangone et al., 2010; Zeiser et al., 2011). However, the Gateway system has disadvantages, including i) ~25 bp *att* recombination sites present between each cassette after assembly, ii) the cost of proprietary enzyme mixes, and iii) the difficulty in assembling more than four cassettes together. In contrast, the SapTrap system i) uses three-base pair junctions, two of which are designed to encode the translation start and STOP codons, ii) is relatively inexpensive, and iii) can efficiently assemble at least 5 cassettes. In principle, the number of cassettes could be increased if desired. The most significant consideration in generating new donor cassette plasmids for SapTrap assembly is that internal SapI sites cannot be present within the donor cassette sequence. Gibson cloning also allows the “scar-free” cloning of transgene vectors, but the specific cloning strategies must be designed for each unique vector. While we have focused on generating transgenes expressed in the hermaphrodite germline, the MosSCI targeting vector pXF87, the gene tag donor cassettes and cloning approach described here should be readily adaptable to expressing transgenes in other tissues.

The advantages of tagging and fluorescently labeling proteins with the HaloTag include increased brightness and photostability (especially compared to red fluorescent proteins) and excellent optical pairing with green fluorescent proteins for 2-color imaging. Additionally, HaloTag labeling offers the flexibility to label a single strain with either JF_549_ HaloTag ligand or JF_646_ HaloTag ligand (Grimm et al., 2015). The disadvantages of HaloTag labeling include the need to introduce the fluorescent ligand (for example, using small scale liquid culture) and the cost of the ligand. Additionally, care should be taken to optimize labeling procedures for each protein to maximize labeling efficiency and minimize background from free ligand. In practice, we find that HaloTag labeling is particularly useful when photobleaching of conventional fluorescent proteins is limiting and/or when imaging in far red is advantageous.

## Materials and Methods

### C. elegans

*C. elegans* hermaphrodite strains were maintained at either 20°C or 25°C on Nematode Growth Medium (NGM) plates containing 3 g/L NaCl, 2.5 g/L peptone and 17 g/L agar supplemented with 1 mM CaCl_2_, 1 mM MgSO_4_, 1 mM KPO_4_ and 5 mg/L Cholesterol with *E. coli* OP50 as a source of food. All strains used in this study are listed in Table 4.

### Cloning

To generate the expression vector pXF87, the two SapI restriction sites in pCFJ350 (Frøkjær-Jensen et al., 2012) were mutated using Q5 Site-Directed Mutagenesis (New England Biolabs) with the oligo pairs XF30F/XF30R and XF31F/XF31R. In addition, the annealed oligos Eg717 and Eg718 were cloned between the XhoI and SpeI sites of pCFJ350.

HaloTag and ceGFP containing PATC-rich endogenous introns were generated in several steps. First, genes were designed *in silico* to minimize germline silencing and increase expression by codon adaptation (Redemann et al., 2011), removal of homology to piRNAs (Batista et al., 2008), and inclusion of a short endogenous intron from *rpl-18* and four synthetic introns (Okkema et al., 1993) using the freely available gene editor ApE (M. Wayne Davis, unpublished). Second, the synthetic genes were synthesized as gBlocks (IDT), cloned into a plasmid, and sequence verified. Third, PATC-rich introns from a gene that is resistant to germline silencing, *smu-1* (Spike et al., 2001), were introduced into the synthetic genes by Golden Gate cloning as described previously (Frøkjær-Jensen et al., 2016). Finally, correct splicing and expression was verified by expression of the synthetic genes with and without PATC-rich introns using an *eft-3* promoter and *tbb-2* 3’UTR.

Donor cassette plasmids numbered pXF, pJF and pSM were generated by cloning PCR products into the pCR BluntII vector backbone using the Zero Blunt™ Topo™ system (Thermo Fisher Scientific). pSDH donor cassette plasmids were cloned by ligating PCR products into pSDH76, a derivative of pCR BluntII containing two XcmI sites that generate T-overhangs following digestion with XcmI. PCR primers for each plasmid are listed in Table 3. pXF87 and all donor plasmids were sequence verified.

**Table 3.**
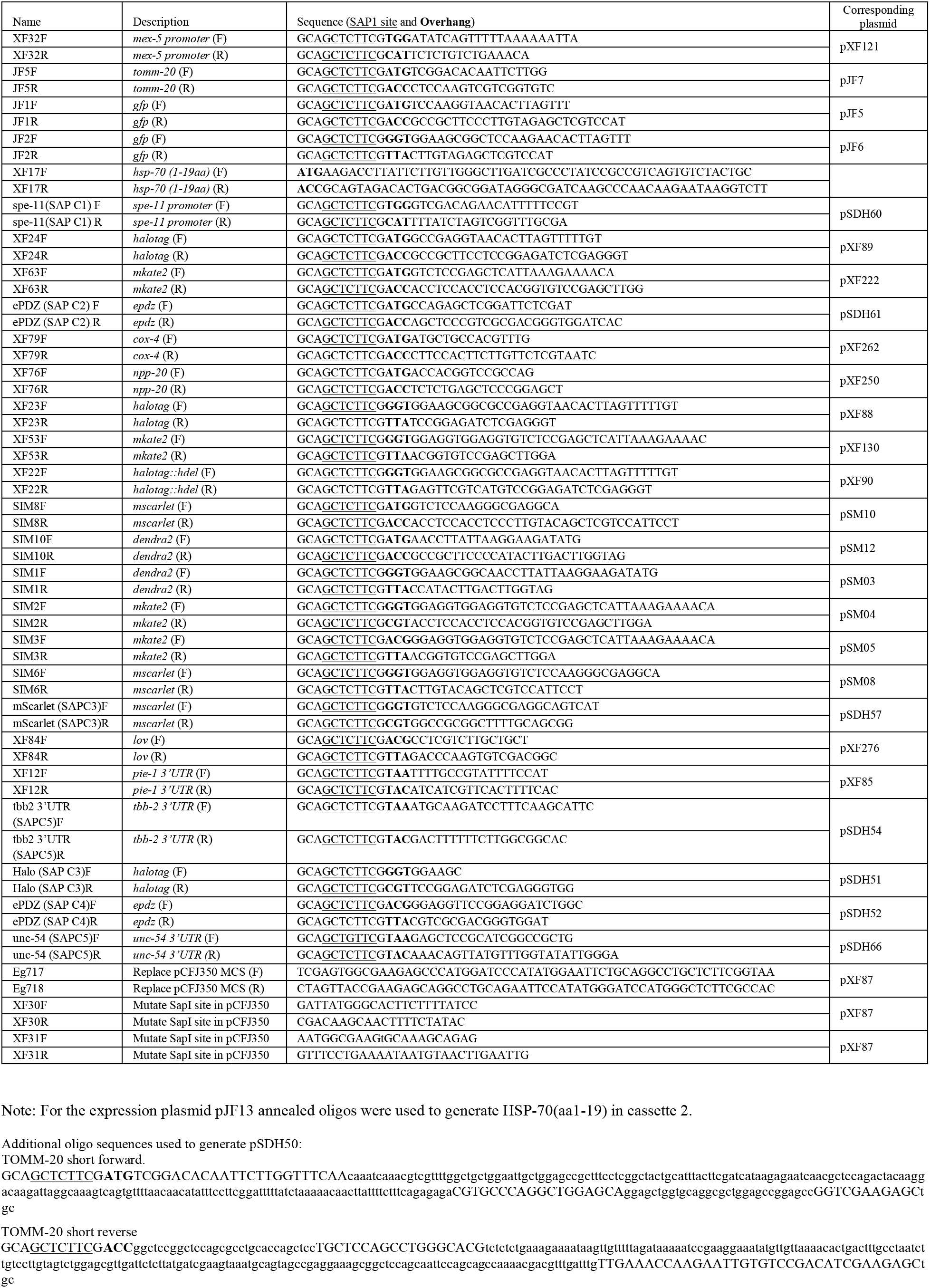
Primers used in this study.

To assemble HSP-70 (aa1-19) into the first cassette position of the expression vector pXF108, 10mM of oligos XF17F and XF17R were gradually cooled from 95°C to 25°C in a BioRad T1000 thermocycler. Annealed oligos were phosphorylated by T4 polynucleotide kinase (NEB) for two hours at 37°C followed by 65°C for 20 minutes. The donor and primers plasmids are listed in Tables 1 and 3, respectively.

**Table 4.** Strains used in this study

| Strain | Genotype | Construction | Reference: |
| --- | --- | --- | --- |
| EG8078 | *oxTi185 I; unc-119(ed3) III* |  | Frokjaer-Jensen et al (2014) |
| EG8079 | *oxTi179 II; unc-119(ed3) III* |  | Frokjaer-Jensen et al (2014) |
| EGD329 | *egxSi126 [unc-119(+), pmex-5::hsp-70(aa1-19)::halotag::HDEL::pie-1 3'UTR] I; unc-119(ed3) III* | Injected pJF13 into EG8078 | This study |
| EGD412 | *egxSi136 [unc-119(+); pmex-5::tomm-20::halotag::pie-1 3'UTR] II; unc-119(ed3) III* | Injected pJF17 into EG8079 | This study |
| EGD496 | *egxSi117 [unc-119 (+); pmex-5::npp-20::gfp::pie-1 3'UTR] I; unc-119(ed3) III* | Injected pXF253 into EG8078 | This study |
| EGD497 | *egxSi118 [unc-119 (+); pmex-5::npp-20::halotag::pie-1 3'UTR] II; unc-119 (ed3) III* | Injected pXF255 into EG8079 | This study |
| EGD549 | *egxSi144 [unc-119 (+); pmex-5::cox-4::halotag::pie-1 3'UTR] II; unc-119 (ed3) III* | Injected pXF266 into EG8079 | This study |
| EGD565 | *egxSi145 [unc-119 (+); pmex-5::hsp-70(aa1-19)::halotag::HDEL::pie-1 3'UTR] II; unc-119 (ed3) III* | Injected pJF13 into EG8079 | This study |
| EGD623 | *egxSi152 [unc-119(+); pmex-5::tomm -20::gfp::pie-1 3'UTR] II; unc-119(ed3) III* | Injected pSM16 into EG8079 | This study |
| EGD629 | *egxSi155 [unc-119(+); pmex-5::tomm -20::mKate2::pie-1 3'UTR] II; unc-119(ed3) III* | Injected pSM20 into EG8079 | This study |
| EGD631 | *egxSi155 [unc-119(+); pmex-5::tomm -20::Dendra2::pie-1 3'UTR] II; unc-119(ed3) III* | Injected pSM17 into EG8079 | This study |
| EGD633 | *egxSi159 [unc-119(+); pmex-5::tomm -20::mScarlet::pie-1 3'UTR] II; unc-119(ed3) III* | Injected pSM22 into EG8079 | This study |
| EGD615 | *cox-4(zu476[cox-4::eGFP::3XFLAG]) I; egxSi136 [unc-119(+); pmex-5::tomm -20::halotag::pie-1 3'UTR] II; unc-119(ed3?) III* | Crossed EGD412 and JJ2586 | This study |
| JJ2586 | *cox-4(zu476[cox-4::eGFP::3XFLAG]) I* |  | Raiders et al, 2018 |
| TBD307 | *dhc-1(he255[epdz::mcherry::dhc-1]) I; utdSi51(Pmex-5::tomm -20(aa1-55)::halotag::lov::tbb-2 3'UTR) II* | Injected pSDH68 into EG8079. Crossed to SV2095. | This study |
| SV2095 | *dhc-1(he255[epdz::mcherry::dhc-1]) I; ruls57[Ppie-1::gfp::tbb-2 + unc119(+)] V* |  | Fielmich et al, 2018 |

### Assembly reaction

Assembly reactions 50 μL included 1 nM of pXF87 and each donor cassette plasmid, 400 units of T4 DNA ligase (NEB), 10 units of SapI enzyme (NEB), 1X NEB CutSmart buffer and 1 mM ATP. For assemblies including annealed oligos, phosphorylated annealed oligos were used at a final concentration of 3 nM in the assembly reaction. Reactions were incubated for 22-24 hours at 25°C, transformed into Stellar Competent cells (Clontech). Four to six plasmid clones were first screened by restriction digest with XhoI and SpeI. Plasmids with the correct restriction digest pattern were sequenced across each cassette boundary. MosSCI targeting vector assembly reactions are listed in Table 2. Note that because the background of unassembled vectors in our assembly reactions was typically low, our protocol omits the counterselection restriction enzyme step described in the original SapTrap protocol (Schwartz and Jorgensen, 2016).

### Transgenesis

Double strand breaks at Mos1 landing sites were generated using CRISPR/Cas9. With the exception of strains EGD615, EGD629, EGD631 and EGD633, injection mixes contained 50 ng/μL assembled MosSCI targeting vectors and pXW7.01 and pXW7.02 sgRNA/Cas9 vectors (gifts from Katya Voronina, University of Montana), which generate double strand breaks at the *ttTi5605* universal MosSCI insertion site. For strains EGD615, EGD629, EGD631 and EGD633, injection mixes contained 0.25 μg/μL Cas9 protein, 0.1μg/μL tracrRNA, 0.028 μg/μL crRNAs BH0278

(GCGUCUUCGTACCUUUUUGGGUUUUAGAGCUAUGCUGUUUUG) and BH0279 (GUCCCAUCGAAGCGAAUAGGGUUUUAGAGCUAUGCUGUUUUG) (Dharmacon) and 0.1 μg/μL assembled MosSCI plasmids. The universal MosSCI strains EG8078 or EG8079 (Frøkjær-Jensen et al., 2014) were injected, singled and incubated for 10 days at 20°C. ~10 worms from plates containing non-Unc animals were transferred to new plates. Plates that stably gave rise to non-*unc* progeny were visually screened for fluorescent transgene expression.

### HaloTag staining

20 to 30 L4 worms were stained in 25 μL S media containing concentrated OP50 bacteria and 2.5 μM of either JF_549_ HaloTag ligand or JF_646_ HaloTag ligand (Grimm et al., 2015) in a darkened 96 well plate shaking at 150 rpm for 19 hours at 23°C. Water was placed in the neighboring wells to help prevent evaporation. Animals were recovered on NGM plates for up to two hours before imaging.

### MitoTracker Deep Red staining

L4 worms were fed overnight on an NGM plate that had been seeded with 100 μL concentrated OP50 bacteria mixed with 1 μL of 1 mM MitoTracker Deep Red FM dye (Cell Signaling Technology, Cat #8778S).

### Imaging

With the exceptions of the TOMM-20::Dendra2 strain and optogenetic strains (Figure 4), all images were collected on a spinning-disk microscope built on a Nikon Eclipse Ti base and equipped with an Andor CSU-W1 two camera spinning disk module, Zyla sCMOS cameras, an Andor ILE laser module and a Nikon 100X Plan Apo 1.45 NA oil immersion objective (Micro Video Instruments, Avon, MA).

TOMM-20::Dendra2 was imaged on a Marianas spinning disk microscope (Intelligent Imaging Innovations) built around a Zeiss Axio Observer Z.1 equipped with a Photometrics Evolve EMCCD camera, 50 mW 488 and 561 nm solid state lasers, a CSU-X1 spinning disk (Yokogawa, Tokyo Japan) and a Zeiss 100X Plan-Apochromat objective. Photoconversion was performed by 5 second illumination with a 405 epifluorescent light.

To stimulate the relocalization of mitochondria (Figure 4), embryos were illuminated with a 50 mW 640 nm solid-state laser used to excite MitoTracker DeepRed (20% laser power, 100 msec exposure, camera gain of 1) and a 50 mW 488 nm solid-state laser used to stimulate the interaction between ePDZ and LOV domains (80% laser power and 100 msec exposure). A Plan-Apochromat 100x/1.4 NA oil immersion DIC objective (Zeiss) was used and Z-stacks (one micron step size, 11 steps) were collected at 60-second intervals. The images displayed in Figure 4 are maximum intensity projections of three Z planes from the cell midplane.

### Data and reagent availability

Strains and reagents are available upon request. Supplemental materials describing the sequence of tag donor cassettes are available through the GSA FigShare portal.

## Acknowledgements

We thank Bing He and members of the Griffin lab for comments on the manuscript. We thank Katya Voronina (U. of Montana) for the plasmids pXW7.01 and pXW7.02. We thank Ann Lavanway and Zdenich Zvindrych of the Dartmouth Life Sciences Imaging Facility for assistance with imaging. The Molecular Biosciences core facility is supported by the Norris Cotton Cancer Center and by NCI grant 5P30CA023108-40. This work was supported by grants from the NIH (R01GM110194 to EEG), baseline funding from KAUST (to CFJ) and the NWO (016.Veni.181.051 to SDH). Erik Jorgensen (U. of Utah, HHMI) contributed resources to generate some of the transgenes from NIH grant R01GM095817.

---

**Figure S1.**
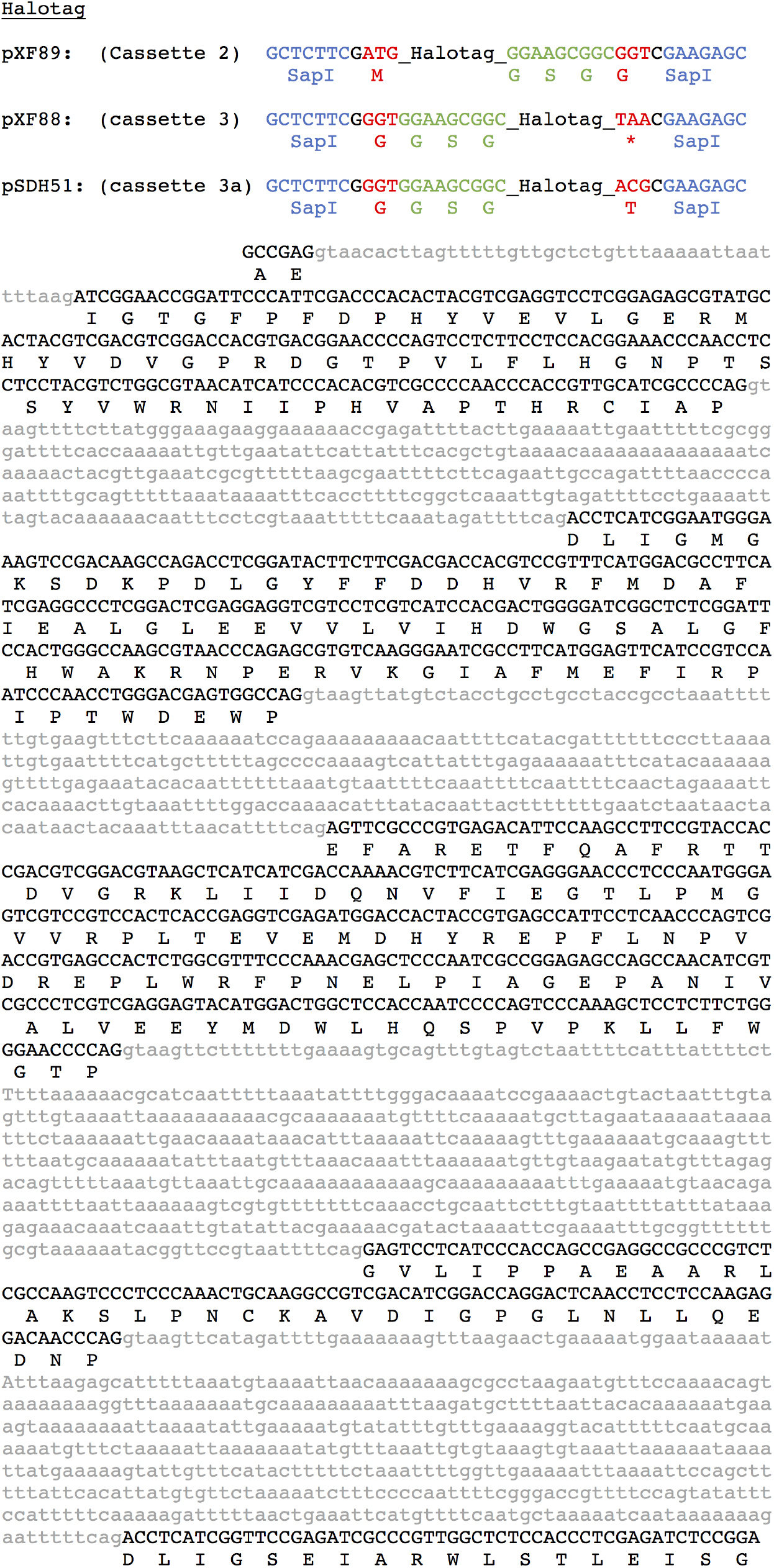
Sequence of HaloTag donor cassettes. Cassette-specific flanking sequences are shown at the top. The SapI recognition sites are in blue and the cleavages sites are in red.

**Figure S2.**
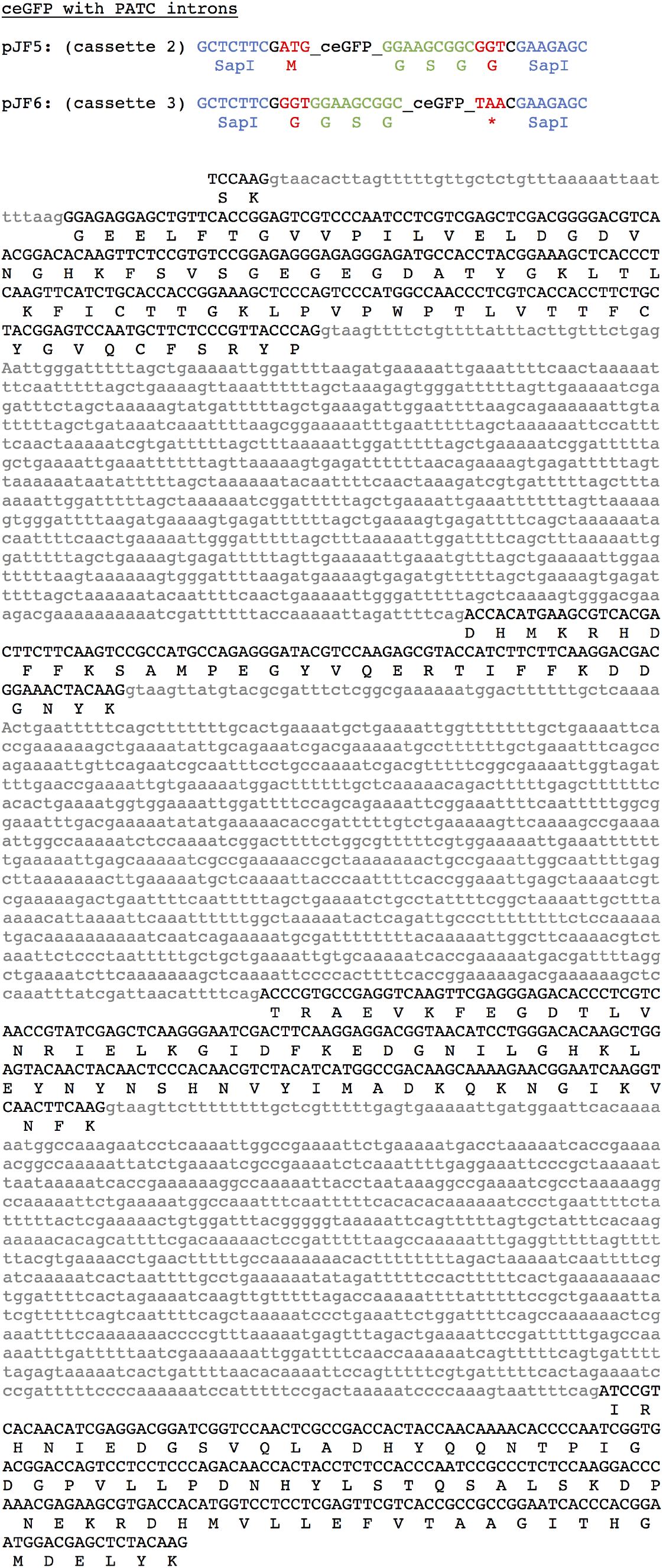
Sequence of ceGFP donor cassettes. Cassette-specific flanking sequences are shown at the top.The SapI recognition sites are in blue and the cleavages sites are in red.

**Figure S3.**
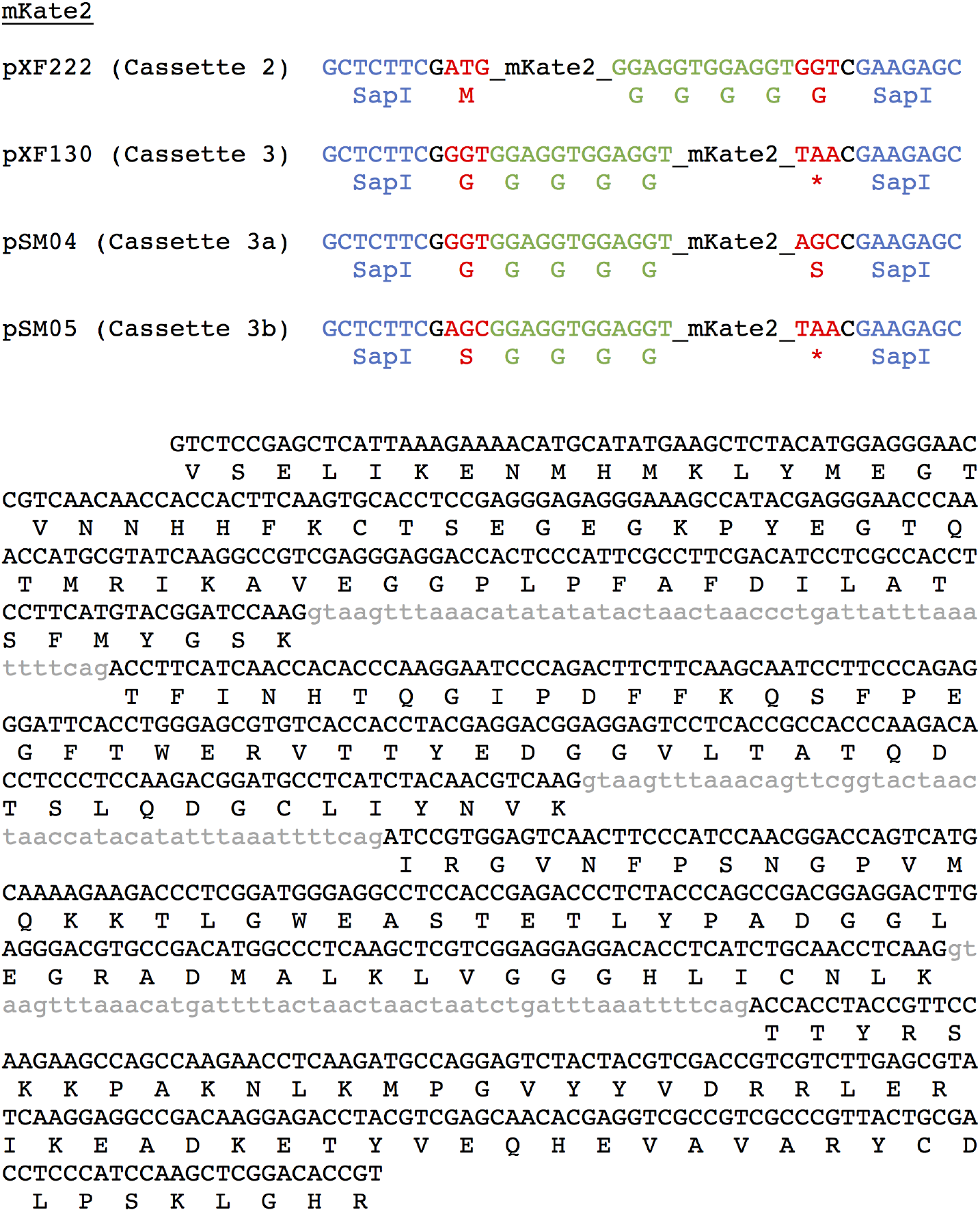
Sequence of mKate2 donor cassettes. Cassette-specific flanking sequences are shown at the top. The SapI recognition sites are in blue and the cleavages sites are in red.

**Figure S4.**
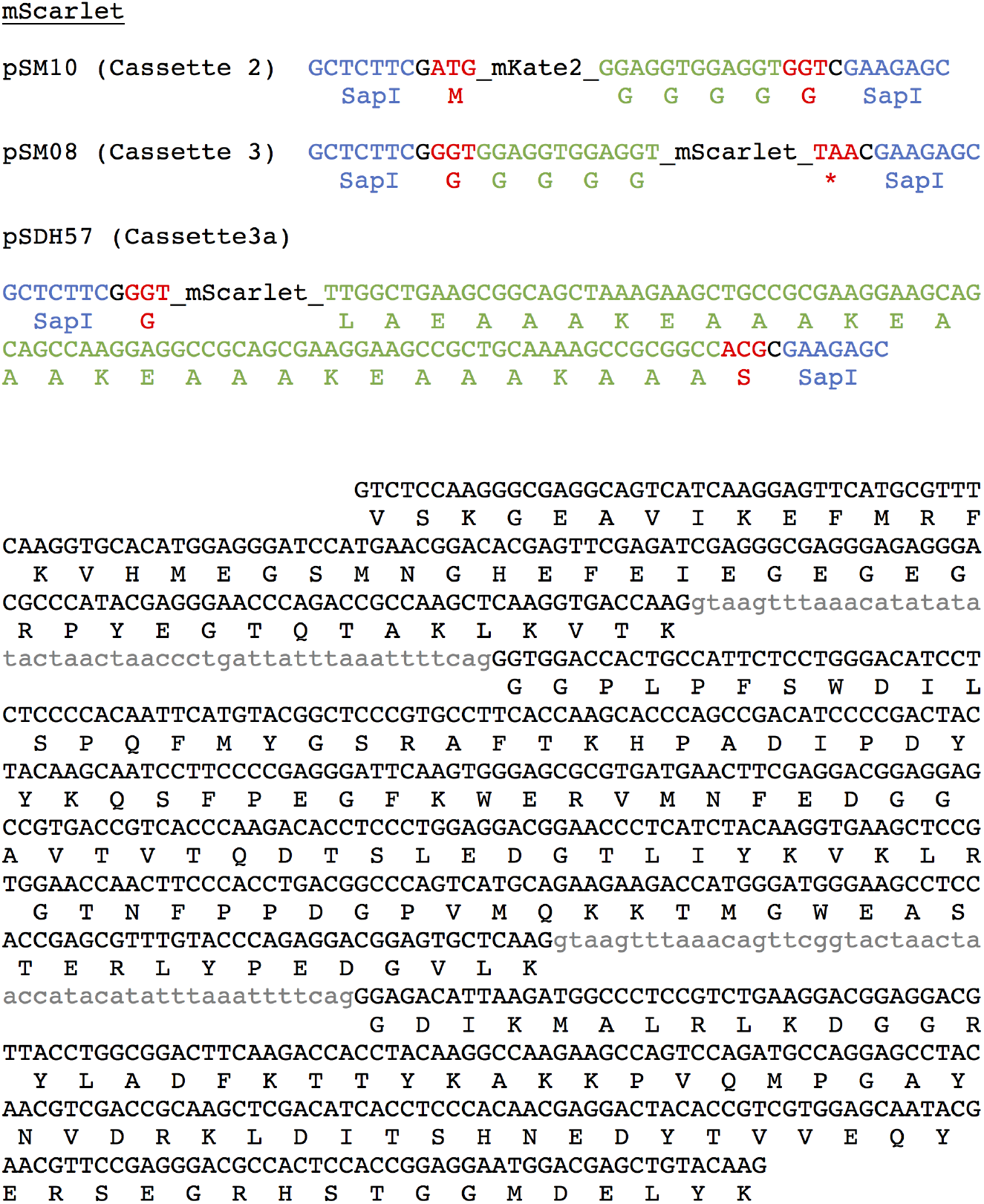
Sequence of mScarlet donor cassettes. Cassette-specific flanking sequences are shown at the top. The SapI recognition sites are in blue and the cleavages sites are in red.

**Figure S5.**
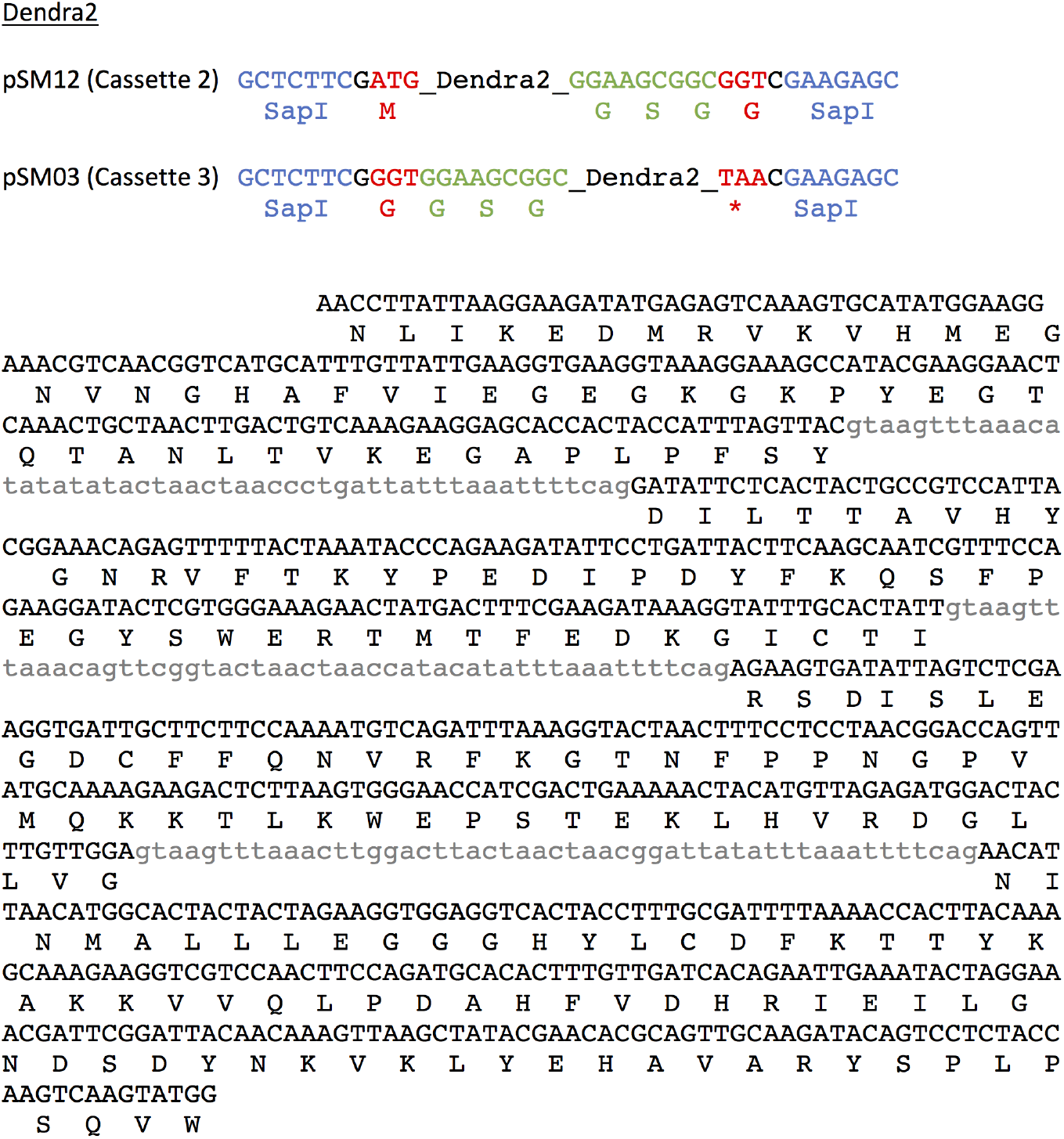
Sequence of Dendra2 donor cassettes. Cassette-specific flanking sequences are shown at the top. The SapI recognition sites are in blue and the cleavages sites are in red.

**Figure S6.**
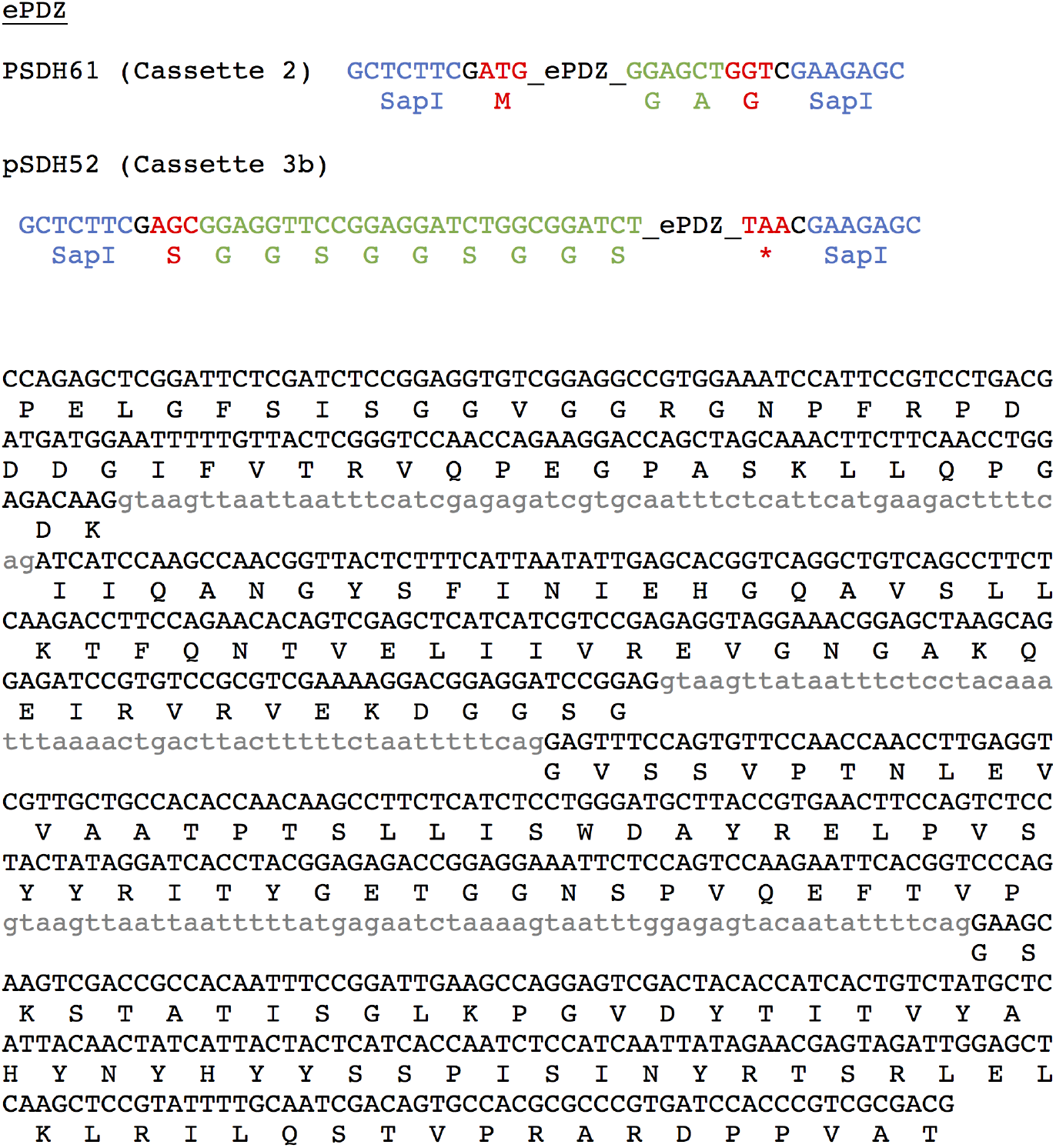
Sequence of ePDZ donor cassettes. Cassette-specific flanking sequences are shown at the top. The SapI recognition sites are in blue and the cleavages sites are in red.

**Figure S7.**
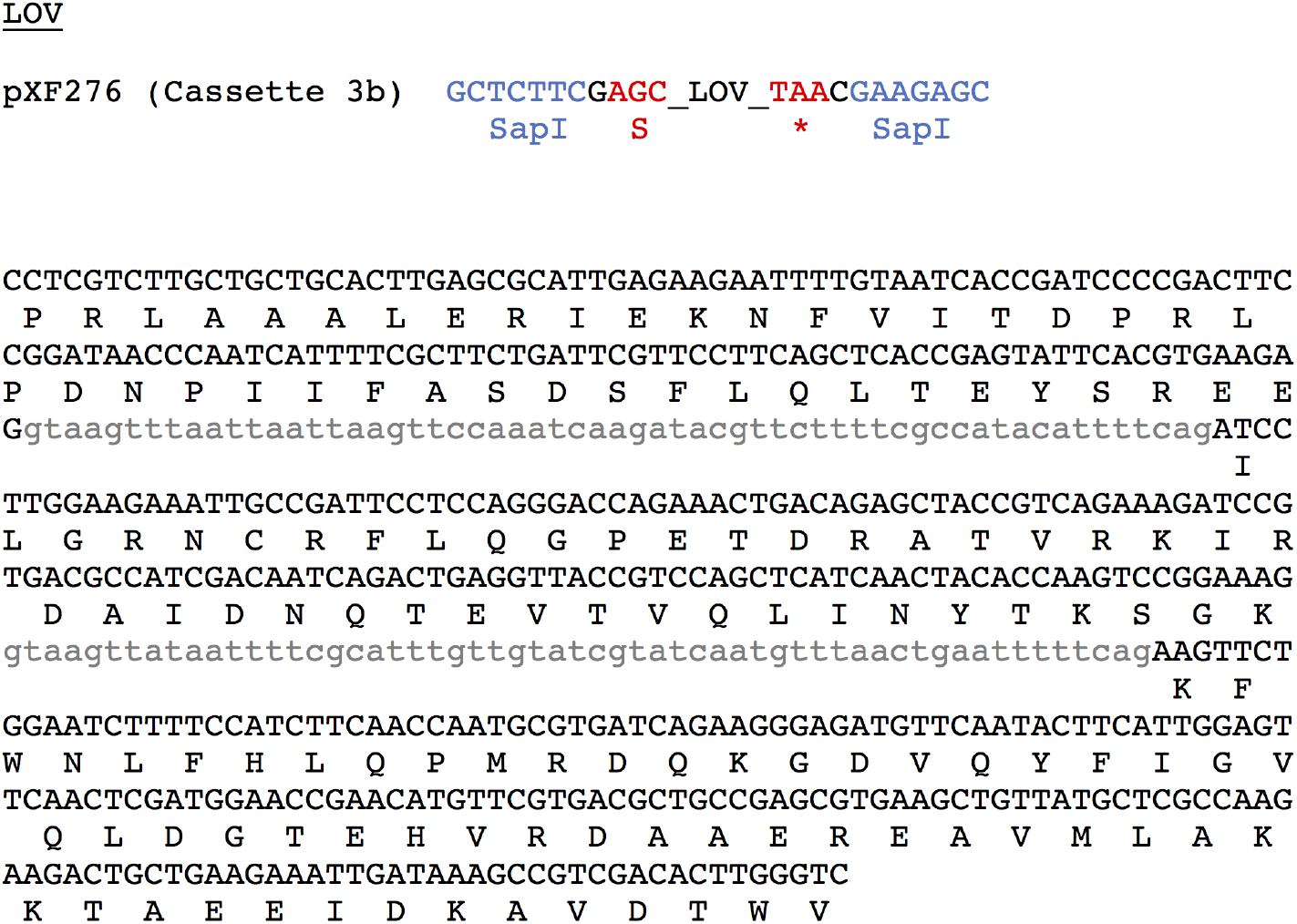
Sequence of LOVdonor cassette. Flanking sequences are shown at the top. The SapI recognition sites are in blue and the cleavages sites are in red.

## Literature Cited

Batista, P. J., J. G. Ruby, J. M. Claycomb, R. Chiang, N. Fahlgren, et al., 2008 PRG-1 and 21U-RNAs interact to form the piRNA complex required for fertility in *C. elegans*. Mol. Cell 31: 67–78.

Brasch, M. A., J. L. Hartley, and M. Vidal, 2004 ORFeome cloning and systems biology: standardized mass production of the parts from the parts-list. Genome Res. 14:2001–2009.

D’Arcangelo, J. G., K. R. Stahmer, and E. A. Miller, 2013 Vesicle-mediated export from the ER: COPII coat function and regulation. Biochim. Biophys. Acta 1833:2464–2472.

Dupuy, D., Q. R. Li, B. Deplancke, M. Boxem, T. Hao, et al., 2004 A first version of the *Caenorhabditis elegans* Promoterome. Genome Res. 14: 2169–2175.

Engler, C., R. Kandzia, and S. Marillonnet, 2008 A one pot, one step, precision cloning method with high throughput capability. PLoS One 3: e3647.

Fielmich, L. E., R. Schmidt, D. J. Dickinson, B. Goldstein, A. Akhmanova, et al., 2018 Optogenetic dissection of mitotic spindle positioning in vivo. Elife 7: e38198.

Frøkjaer-Jensen, C., M. W. Davis, M. Ailion, and E. M. Jorgensen, 2012 Improved Mos1-mediated transgenesis in *C. elegans*. Nat. Methods 9: 117–118.

Frøkjaer-Jensen, C., M. W. Davis, C. E. Hopkins, B. J. Newman, J. M. Thummel, et al., 2008 Single-copy insertion of transgenes in *Caenorhabditis elegans*. Nat. Genet. 40: 1375–1383.

Frøkjaer-Jensen, C., M. W. Davis, M. Sarov, J. Taylor, S. Flibotte, et al., 2014 Random and targeted transgene insertion in *Caenorhabditis elegans* using a modified Mos1 transposon. Nat. Methods 11: 529–534.

Frøkjaer-Jensen, C., N. Jain, L. Hansen, M. W. Davis, Y. Li, et al., 2016 An Abundant Class of Non-coding DNA Can Prevent Stochastic Gene Silencing in the *C. elegans* Germline. Cell 166: 343–357.

Gibson, D. G., L. Young, R. Y. Chuang, J. C. Venter, C. A. Hutchison 3rd, et al., 2009 Enzymatic assembly of DNA molecules up to several hundred kilobases. Nat. Methods 6: 343–345.

Grimm, J. B., B. P. English, J. Chen, J. P. Slaughter, Z. Zhang, et al., 2015 A general method to improve fluorophores for live-cell and single-molecule microscopy. Nat. Methods 12: 244–250.

Hartley, J. L., G. F. Temple, and M. A. Brasch, 2000 DNA cloning using in vitro site-specific recombination. Genome Res. 10: 1788–1795.

Kelly, W. G., S. Xu, M. K. Montgomery, and A. Fire, 1997 Distinct requirements for somatic and germline expression of a generally expressed *Caenorhabditis elegans* gene. Genetics 146: 227–238.

Mangone, M., A. P. Manoharan, D. Thierry-Mieg, J. Thierry-Mieg, T. Han, et al., 2010 The landscape of *C. elegans* 3’UTRs. Science 329: 432–435.

Merritt, C., D. Rasoloson, D. Ko, and G. Seydoux, 2008 3’ UTRs are the primary regulators of gene expression in the *C. elegans* germline. Curr. Biol. 18: 1476–1482.

Nance, J., and C. Frøkjær-Jensen. (2019). The *Caenorhabditis elegans* Transgenic Toolbox. Genetics 212, 959–990.

Okkema, P. G., S. W. Harrison, V. Plunger, A. Aryana, and A. Fire, 1993 Sequence requirements for myosin gene expression and regulation in *Caenorhabditis elegans*. Genetics 135: 385–404.

Philip, N. S., F. Escobedo, L. L. Bahr, B. J. Berry, and A. P. Wojtovich, 2019 Mos1 Element-Mediated CRISPR Integration of Transgenes in *Caenorhabditis elegans*. G3 9: 2629–2635.

Raiders, S.A., M. D. Eastwood, M. Bacher, and J. R. Priess, 2018 Binucleate germ cells in *Caenorhabditis elegans* are removed by physiological apoptosis. PLoS Genet. 14: e1007417.

Redemann, S., S. Schloissnig, S. Ernst, A. Pozniakowsky, S. Ayloo, et al., 2011 Codon adaptation-based control of protein expression in *C. elegans*. Nat. Methods 8: 250–252.

Schwartz, M. L., and E. M. Jorgensen 2016 SapTrap, a Toolkit for High-Throughput CRISPR/Cas9 Gene Modification in *Caenorhabditis elegans*. Genetics 202: 1277–1288.

Seth, M., M. Shirayama, W. Tang, E. Z. Shen, S. Tu, et al., 2018 The Coding Regions of Germline mRNAs Confer Sensitivity to Argonaute Regulation in C. elegans. Cell Rep. 22: 2254–2264.

Shirayama, M., M. Seth, H. C. Lee, W. Gu, T. Ishidate, et al., 2012 piRNAs initiate an epigenetic memory of nonself RNA in the *C. elegans* germline. Cell 150: 65–77.

Siniossoglou, S., C. Wimmer, M. Rieger, V. Doye, H. Tekotte, et al., 1996 A novel complex of nucleoporins, which includes Sec13p and a Sec13p homolog, is essential for normal nuclear pores. Cell 84: 265–275.

Spike, C. A., J. E. Shaw, and R. K. Herman, 2001 Analysis of *smu-1*, a gene that regulates the alternative splicing of unc-52 pre-mRNA in *Caenorhabditis elegans*. Mol. Cell Biol. 21: 4985–4995.

Strickland, D., Y. Lin, E. Wagner, C. M. Hope, J. Zayner, et al., 2012 TULIPs: tunable, light-controlled interacting protein tags for cell biology. Nat Methods 9: 379–384.

Zeiser, E., C. Frøkjaer-Jensen, E. Jorgensen, and J. Ahringer, 2011 MosSCI and gateway compatible plasmid toolkit for constitutive and inducible expression of transgenes in the C. elegans germline. PLoS One 6: e20082.

Zhang, D., S. Tu, M. Stubna, W. S. Wu, W. C. Huang et al., 2018 The piRNA targeting rules and the resistance to piRNA silencing in endogenous genes. Science 359: 587–592.

